# Inhibition of Yes-Associated Protein (YAP) with Verteporfin Enhances Radiosensitivity in Chordoma by Inducing G_2_M Arrest and Inhibiting the DNA Damage Response

**DOI:** 10.1101/2024.11.03.621728

**Authors:** Oluwaseun O. Akinduro, Paola Suarez-Meade, Paula Schiapparelli, Rachel Whitehead, Stephany Y. Tzeng, Steven S. Rosenfeld, Sungjune Kim, Claudia Palena, Ziya L. Gokaslan, Jordan J. Green, Alfredo Quinones-Hinojosa

## Abstract

Chordomas are locally invasive cancers that are highly resistant to radiotherapy. The Brachyury and Yes-Associated Protein (YAP) regulatory axis has been implicated as the primary driver of tumorigenicity in chordoma. Here, we aimed to enhance chordoma radiosensitivity by repurposing the FDA-approved YAP inhibitor, Verteporfin. We used five patient-derived chordoma cell lines and generated two YAP1 knockdown cell lines to validate the YAP-targeting phenotype in chordoma. Verteporfin treatment reduced the expression of DNA damage repair proteins and genes. YAP inhibition with either verteporfin or YAP knockdown resulted in enhanced DNA double-stranded breaks after radiation via inhibition of the DNA damage repair pathway and accumulation of cells in the G_2_M phase. Verteporfin inhibited chordoma tumor growth alone and in combination with radiation in a xenograft mouse model treated with verteporfin loaded microparticles, resulting in sensitization of chordoma tumors to radiation. YAP inhibition with verteporfin renders chordoma more sensitive to radiation via inhibition of the DNA damage repair cascade and accumulation of cells in G_2_M when they are most susceptible to radiation damage.

## INTRODUCTION

Chordomas are locally aggressive bone tumors that arise from notochordal remnants and have a very poor prognosis because of their high recurrence rate and invasion of local structures. ^1-4^ They are the most common malignant primary bone tumors and have a predilection for growth in the midline axial skeleton, most commonly in the clivus and sacrum. Chordomas have a ten-year survival of approximately 41%, are highly recurrent and radioresistant, and have no FDA-approved targeted therapies. ^5,2^ Their midline growth patterns cause them to invade critical neural, vascular, and bone structures, making surgical resection of chordoma very challenging. ^6^ When feasible, the gold standard treatment for these tumors is *en-bloc* resection with wide margins; if tumor-free margins cannot be obtained, surgery should be followed by high-dose proton-based radiation therapy. ^7^ The presence of the brainstem, spinal cord, cranial nerves, and pelvic organs limits the safety of high-dose radiation, and patients may develop debilitating radiation-toxicity complications. Several chemotherapies and targeted treatments have been tested for the treatment of chordomas but have failed to prevent tumor recurrence and metastasis. ^8,9,10,11^ Therefore, effective adjuvant therapy is urgently needed to increase the radiation sensitivity of chordomas to lengthen progression-free survival times and improve patient prognosis.

The embryonic transcription factor brachyury drives tumorigenicity in all chordomas, making it an attractive therapeutic target. ^12^ Brachyury promotes tumor progression and drives stemness in chordoma through the activation of yes-associated protein (YAP), which is one of the main components of the Hippo pathway. ^13,14^ YAP associates with its paralog TAZ (transcriptional co-activator with PDZ-binding motif) and translocates to the nucleus to bind to the TEA domain (TEAD) transcription factor and exert its downstream tumorigenic effects. ^14^ Brachyury driven-YAP activity can be inhibited with the use of the FDA-approved drug verteporfin. ^15,16^ Verteporfin prevents YAP-TEAD interactions, inhibits YAP transcriptional activity, and thus impedes tumor progression.^17^

In this study, we aimed to optimize radiation therapy in chordomas and to understand the mechanisms underlying chordoma radioresistance. We hypothesized that the inhibition of YAP with verteporfin would render chordomas more sensitive to radiation therapy via inhibition of the DNA damage response pathway and modulation of the cell cycle (**Figure 1**). We used novel microparticles containing verteporfin designed by our group and previously tested them in other types of cancer. ^15,16,18^ These verteporfin-loaded microparticles allow the sustained release of the drug over time. This would be expected to prolong the exposure time of the tumor to this drug, which has a short half-life (6 h) when administered via conventional intravenous means. In this study, we present the first in-depth examination of the mechanism of verteporfin-induced radiation sensitivity in chordoma.

**Figure 1.**
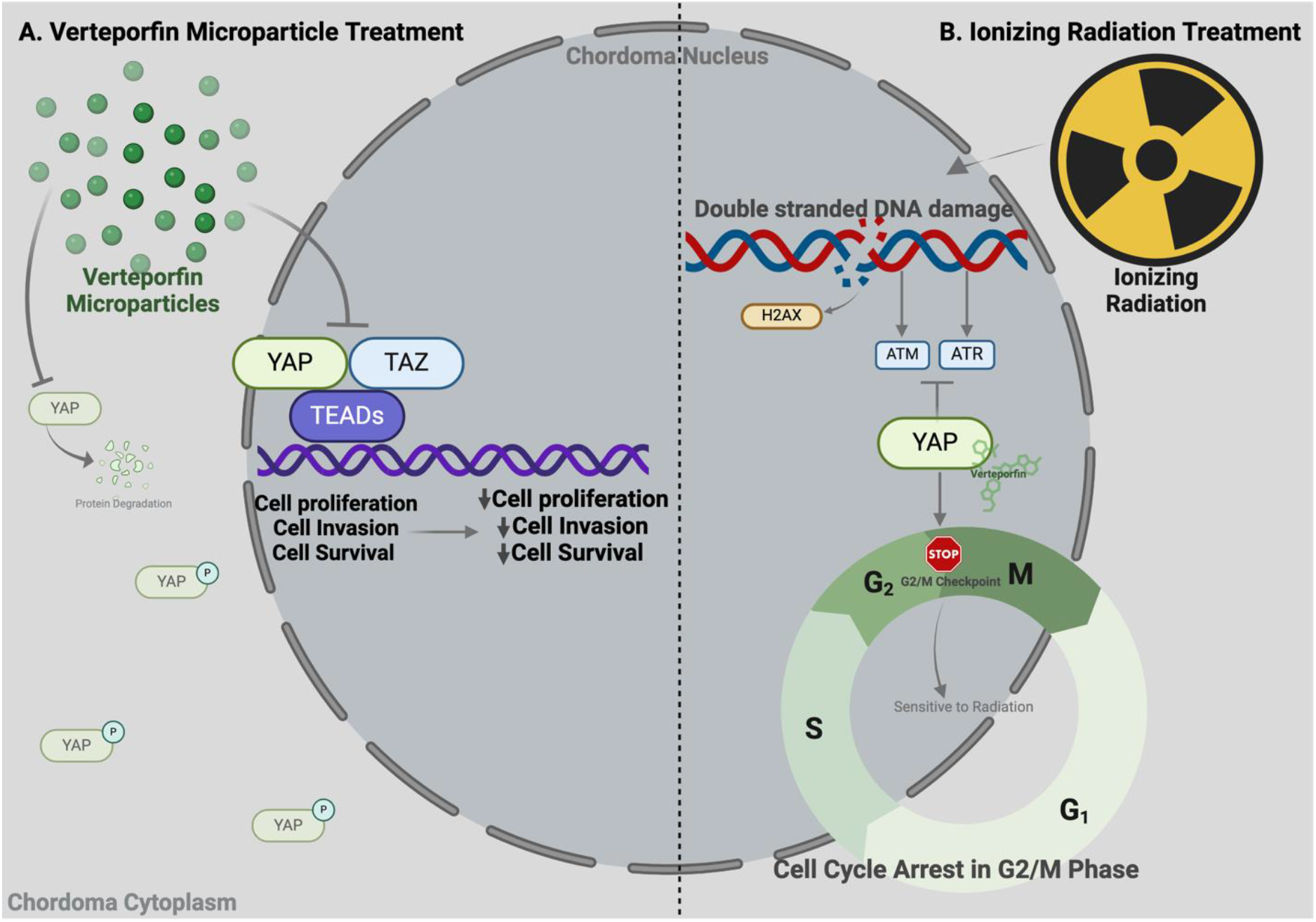
Graphical representation of Verteporfin Mechanism in Chordoma.**A.**Brachyury is one of the main drivers of tumorigenesis in chordomas; this transcriptional factor is present in all chordoma tumors and has been demonstrated to increase the expression of the Yes-associated protein (YAP), a oncoprotein located in the cytoplasm in an inactive form. When YAP is activated by brachyury, it translocates to the nucleus to bind to TEAD, for the activation of genes responsible for proliferation, migration, and apoptosis. **B**. Verteporfin is an FDA approved drug that prevents YAP from binding to the TEAD transcription factor. **C**. Ionizing radiation causes double stranded DNA damage which activates the DNA damage repair cascade including γH2AX, ATM, and ATR. Verteporfin inhibits YAP, which modulates the cell cycle and inhibits the DNA damage repair process.

## METHODS

### Cell Culture

The chordoma cell lines JHC7, U-CH1, U-CH2, UM-Chor1, and MUG-Chor1 were used in this study. Cell line JHC7 was established from patient-derived tissues and cultured as previously described.^19,20^ U-CH1 (CLR-3217), U-CH2 (CLR-3218), UM-Chor 1 (CRL-3270), and MUG-Chor1 (CRL-3219) cell lines were purchased from ATCC and cultured in MesenPRO basal medium with an associated growth supplement (Invitrogen), GlutaMAX (Invitrogen), and 5% antibiotic/antimycotic (Invitrogen).

### Virus Preparation and Viral Transduction for YAP Knockdown

Glycerol stocks were obtained from the Human Mission TRC1 sequence-verified shRNA lentiviral plasmid vectors. Plasmids were isolated using a Spin Miniprep Kit (Qiagen) and verified via enzymatic restriction. Lentiviral particles were produced by co-transfecting 293T cells with Lipofectamine 3000 (Invitrogen), an shRNA transducing vector, and two packaging vectors, psPAX2 and pMD2.G. On day 3 after transfection, media containing the virus were harvested and filtered through a 0.22 mm pore cellulose acetate filter and then concentrated with the Lenti-X Concentrator (Takara Bio) before quantitative characterization with the Lenti-X p24 Rapid Titer Kit (Takara Bio). Empty TRC1 lentiviral vectors were used as control viruses. JHC7 and UCH1 cells were transduced with equal titers of the virus in growth media supplemented with polybrene (Sigma). 48 hours after transduction, the cells were cultured in complete chordoma media for 24 h prior to selection with puromycin (1 mg/mL, Sigma).

### Verteporfin Treatment in Vitro

Verteporfin (Sigma-Aldrich) stock solution was solubilized in dimethyl sulfoxide (DMSO) at 2 mg/mL to generate a stock concentration of 2.78mM, as per the manufacturer’s recommendation. Cells were seeded and grown until they reached approximately 60-80% confluency prior to verteporfin treatment. *In vitro* treatment solutions were prepared by diluting concentrated verteporfin in chordoma cell culture media to the appropriate concentrations for each experimental condition. All treatments were performed using a sterile technique under dark conditions to avoid light exposure to verteporfin.

### In vitro Cell Radiation

Cells were plated in a 2D monolayer under sterile conditions. 24 hours after plating, cells were treated with single doses of radiation (Grays) according to the experimental conditions, using the xRad160 irradiator (Precision X-Ray). A dose-response curve was constructed to identify the *in vitro* IC50 radiation dose for use in subsequent experiments. For experiments that required pretreatment with verteporfin, cells were irradiated 24 h after verteporfin treatment.

### Western Blot

Cells were treated after reaching 60-80% confluency and then processed for western blotting following standard procedures using total protein lysates. Cytoplasmic proteins were obtained using a lysis buffer containing 50 mmol/L HEPES (pH 7.5), 150 mmol/L NaCl, 1% Triton X-100, 1.5 mmol/L MgCl2, EGTA, 10 mmol/L (pH 7.5), 10% glycerol, and inhibitors (0.1 mmol/L Na3VO4, 1% phenylmethylsulfonyl fluoride, and 20 mg/mL aprotinin). After collection of cytoplasmic proteins, the nuclei were lysed with a nuclear buffer containing 20 mmol/L HEPES (pH 8), 0.1 mmol/L EDTA, 5 mmol/L MgCl2, 0.5 mol/L NaCl, 20% glycerol, 1% Nonidet P40, inhibitors (as above), and then quantified. The following primary antibodies were used: anti-YAP (Rabbit Ab, cat 4912S, Cell Signaling), Anti-phosphorylated Yap (pYAP) (Rabbit Ab, cat 4911A, Cell Signaling, S127), anti-ATM (Rabbit Ab, cat 2873, Cell Signaling), anti-ATR (Rabbit Ab, cat 3767, Biovision), anti-cABL (Rabbit Ab, cat 2862, Cell Signaling), anti-PARP (Rabbit Ab, cat 3002 Ab, Biovision), and Anti-Phospho-Histone H2A.X (Ser139) (Mouse Ab, Clone JBW301, Millipore Sigma). Normalization was performed using β-actin or alpha-tubulin staining. Three replicate gels were used to obtain p-values. The relative pixel intensity was calculated using Fiji Image J.

### Quantitative PCR (qPCR)

Total RNA was isolated by adding TRIzol Reagent (Thermo Fisher Scientific -Life Technologies) directly to the cells. Total RNA was extracted using an RNeasy Mini Kit (Qiagen, Hilden, Germany). Total RNA from each sample was reverse transcribed into complementary DNA (cDNA). cDNA was synthesized using primed reverse transcriptase (Thermo Fisher Scientific - LifeTechnologies). Real-time qPCR analyses were carried out with triplicate samples of each cDNA sample on a QuantStudio 3 Real-Time PCR system (Applied Biosystems) with SYBR Green Master Mix (Applied Biosystems). The expression levels were calculated relative to GAPDH expression levels. Primer sequences are listed in Supplementary Table 1.

### Colony Formation Assays

The cells were placed in 6-well plates and allowed to attach overnight. JHC7 cells were seeded at 1000 cells/well and treated with verteporfin 24 h after plating. The cells were then irradiated 24 h after verteporfin treatment and allowed to grow for 21 days. Finally, the medium was removed, the cells were fixed with 4% paraformaldehyde, and the colonies were stained with 2.3% crystal violet and counted using a BioSpectrum 810 Imaging System. Cells were considered to have formed colonies when there were at least 50 cells in each group.

### Flow Cytometry Cell Cycle Analysis

Cell cycle analysis was performed using the Click-iT™ EdU Alexa Fluor™ 647 Flow Cytometry Assay Kit (cat. C10424; Invitrogen) according to the manufacturer’s instructions. Cells were analyzed using a CytoFLEX S Beckman Coulter benchtop flow cytometer. The analysis was based on 10,000 singlets in triplicate using the CyExpert software (Beckman Coulter).

### Flow Cytometry H2AX Analysis

Cells were counted, harvested, and stained using the viability dye LIVE/DEAD™ Fixable Aqua Dead Cell Stain Kit (Invitrogen™, cat L34957). After fixation with 4% PFA, the cells were stained using FITC anti-H2A.X Phospho (Ser139) antibody (BioLegend, cat 613403). Cells were analyzed using a CytoFLEX S Beckman Coulter benchtop flow cytometer. The analysis was based on 10,000 singlets in triplicate using the CyExpert software (Beckman Coulter).

### Comet Assay

To analyze double-stranded DNA (dsDNA) damage in cells after verteporfin and radiation treatment, microscope slides were covered with NMP agarose (1.5% v/v), and then chordoma cells (1×10^4^ cells) were embedded in low melting-point agarose and placed on the covered slides. After cell lysis with Triton X-100 in the dark, slides were placed in an alkaline electrophoresis solution (30mM NaOH and 1mM Na_2_EDTA, pH>13) to unwind DNA strands and remove histone proteins. Electrophoresis was performed for 30 min under cold conditions. Slides were then washed with neutralization buffer 40mM Tris-HCl, Ph 7.5). The slides were stained with ethidium bromide and analyzed using a confocal microscope (Zeiss LSM 800). Confocal images were exported and analyzed using Fiji Image J (2020).

### Microparticle Formulation and Characterization

For the microparticle formulation, polylactic-co-glycolic acid (PLGA) with a 85:15 ratio of lactide-to-glycolide was used (molecular weight of 190-240kDa) (ResomerÒ RG 858S, Evonik). PLGA (100 mg) was dissolved in 2mL of dichloromethane (DCM), and this solution was mixed with 1mL of 5 mg/mL verteporfin in DMSO. The solution was homogenized in 50mL of 1% poly(vinyl alcohol) (PVA, 25kDa, 88% hydrolyzed) at 15 krpm for 1 min (Ika T25 digital Ultra-Turrax) to form an emulsion. The emulsion was stirred continuously for 4h to allow solvent evaporation. The resulting microparticles (MPs) were pelleted by centrifugation at 3200 rpm for 5 min and washed four times with diH_2_O. MPs were then lyophilized for 48h and stored as a dry powder at -20°C with desiccant until use. Empty microparticles were fabricated as controls by mixing the PLGA/DCM solution with DMSO only.

### In Vivo Chordoma Injection

All animal protocols were approved by the Institutional Animal Care and Use Committee. Fifty 6-week-old female NOD/SCID mice (Jackson Laboratories, Cat: 005557) were subcutaneously injected in the left flank containing 1 million UCH1 cells that had been previously transduced with lentiviral particles to constitutively express the GFP-Luciferase gene. Before injection, the cells were suspended in a 1:1 mixture of MesenPRO basal medium and Matrigel matrix (Corning). Tumor growth was measured weekly using bioluminescence and caliper measurements. Tumors were allowed to grow to approximately 200 cm3, and then mice were divided into three groups of 16 as follows: 1) vehicle, 2) systemic intraperitoneal free verteporfin, and 3) intratumoral verteporfin-loaded microparticles. Each group of 16 mice was then split so that 8 mice received radiation and 8 mice received sham radiation.

The animals were weighed weekly and continuously evaluated for signs of toxicity. Tumor volume measured with calipers was calculated using the modified ellipsoid formula: tumor volume = ½ (length × width^2^). The individual measuring and calculating the tumor volumes were blinded to the treatments. The animals were euthanized when the tumors had ulcerations or when the tumors met the size/weight limit of our IACUC protocol as follows: 1) tumor size must not exceed 2.0 cm in any direction in an adult mouse, and 2) tumor burden must not be greater than 10% of the animals’ normal body weight.

### In Vivo Verteporfin Treatment

Mice that received intraperitoneal (IP) VP were treated with 100 mg/kg of naked verteporfin (Ambeed, Catalog No. A138297) or 10%:90% DMSO/phosphate-buffered saline solution (IP controls) daily for 14 days. Animals that received intratumoral (IT) verteporfin were treated with two injections of 50 ml of either verteporfin microparticles (containing approximately 320 mg of verteporfin) or sham empty PLGA microparticles on days 1 and 7.

### In Vivo Radiation

Mice were irradiated beginning on day 8 using the Small Animal Image Guided Irradiation System (x-RAD SMART) from Precision X-Ray Irradiation using techniques previously described by our team. ^15,21^ This equipment provides specific targeting accuracy using a fully integrated computed tomography and bioluminescence platform. Mice in the radiation groups received radiation fractions of 5 Gy on each of five consecutive days of 25 Gy for five consecutive days. The animals were monitored daily for signs of radiation side effects or radiation-related skin injuries. Animals that received sham radiation were placed in the irradiator, but no radiation was administered.

### Histology Analysis

At the study endpoint, the mice were euthanized with carbon dioxide, followed by immediate tumor extraction for processing. Tumors were formalin-fixed, paraffin-embedded, and cut into 5 mm sections. Tissue sections were stained with hematoxylin and eosin. Immunohistochemistry was performed using anti-YAP and anti-brachyury antibodies. The slides were scanned using an Aperio AT2 DX System. The percentage of cytoplasmic and nuclear YAP intensity was measured using the Aperio ImageScope x64 Software (Aperio Systems). The signal intensity ranged from 0+ (the lowest detected signal) to 3+ (the strongest detected signal).

### Statistical Analysis and Scientific Rigor

All statistical analyses were performed using the GraphPad Prism Software (version 9.1.0). Analyses were performed using unpaired t-test, one-way analysis of variance (ANOVA) followed by Dunnett’s multiple comparisons test, two-way ANOVA followed by Šídák’s post-hoc multiple comparisons test, and non-linear regression with sum-of-squares F-test. Analyses were performed with a statistical power of 80% and α =0.05. The synergy was calculated using SynergyFinder Plus. ^22^ All *in vitro* experiments were performed in triplicate and repeated three times for scientific rigor.

## RESULTS

### Verteporfin has an additive therapeutic effect when combined with radiation

Our previous work has shown that YAP is a key driver of chordoma development and that it can be targeted by the FDA-approved drug verteporfin. ^11,12^ Hence, we investigated the effect of this drug on radiosensitivity by testing the clonogenicity and dose-response effects of verteporfin treatment in combination with radiation *in vitro*. Verteporfin inhibited chordoma clonogenicity with (2 Gy: **p=0.0439**) and without (0 Gy: p=0.0629) radiation (**Figure 2A)**. We found significantly decreased survival of chordoma cells compared to that of the control (0 Gy, no radiation) (2 Gy: **p<0.0001**, 4 Gy: **p<0.0001**) (**Figure 2B)**. Next, we performed a synergy analysis and found that verteporfin enhanced the effectiveness of radiation when the two were combined, resulting in a mean Bliss score of 5.01, mean Loewe score of 6.99, and mean HSA score of 5.91, indicating that verteporfin and radiation had an additive effect (**Figure 2C**). The combination of 0.25mM of verteporfin and 4 Gy radiation had an HSA synergy score of 26.43, indicating that the combination of the two is synergistic at certain doses. For reference, a synergy score less than -10 is considered antagonistic, a synergy score of -10 to 10 is considered additive, and a synergy score greater than 10 is considered synergistic.

**Figure 2.**
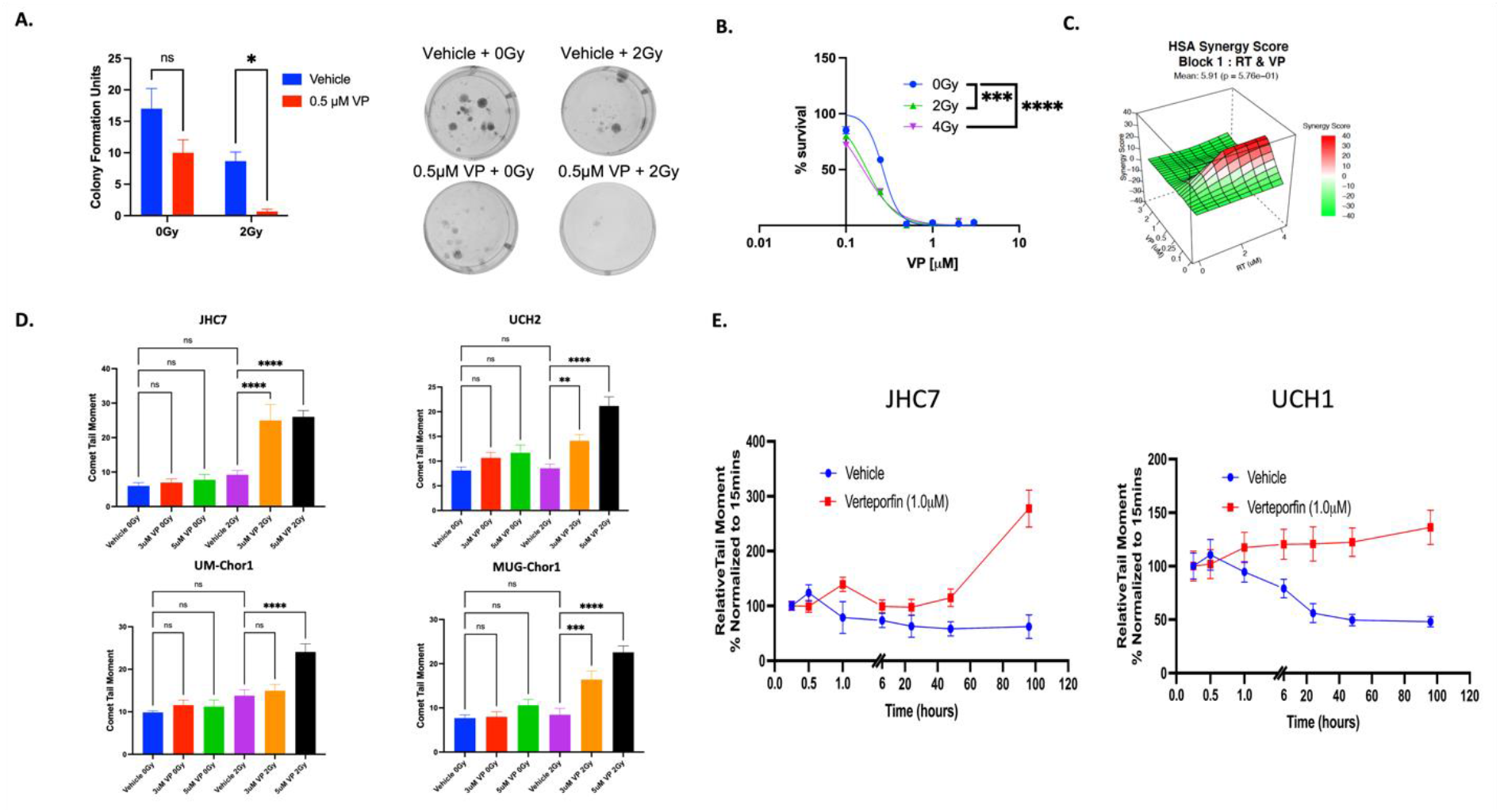
**A**. Verteporfin has an additive effect when combined with radiation. Verteporfin inhibits chordoma clonogenicity with (2Gy: **p=0.0439**) and without (0Gy: p=0.0629) radiation. **B**. Non-linear fit of normalized values demonstrates that increasing doses of verteporfin combined with increasing doses of radiation results in significantly diminished chordoma cell survival (0Gy vs 2Gy: **p=0.0011**, 0Gy vs 4Gy: **p<0.0001**). Non-linear fit extra sum-of-squares F-test. **C**. The combination of verteporfin and radiation results in synergy at certain dose combinations. The combination of 0.25mM of verteporfin and 4Gy radiation had an HSA synergy score of 26.43 indicating high synergy. **D**. The combination of verteporfin and radiation results in significantly increased double-stranded DNA (dsDNA) breaks when compared to vehicle for JHC7 (3*μ*M: **p<0.0001**, 5*μ*M: **p<0.0001**), UCH2 (3μM: **p=0.0042**, 5μM: **p<0.0001**), UM-Chor1 (5μM: **p<0.0001**), and MUG-Chor1 (3μM: **p=0.0009**, 5μM: **p<0.0001**). One-way anova with Šídák’s post hoc analyses. **E**. JHC7 and UCH1 chordoma cells repair dsDNA damage over time but verteporfin prevents DNA repair with significantly increased dsDNA damage at 1hr (JHC7: **p<0.0001**, UCH1: **p=0.0003**), 6hrs (JHC7: **p<0.0001**, UCH1: **p<0.0001**), 24hrs (JHC7: **p=0.0059**, UCH1: **p<0.0001**), 48hrs (JHC7: **p<0.0001**, UCH1: **p<0.0001**), and 96hrs (JHC7: **p<0.0001**, UCH1: **p<0.0001**). Two-way anova with Šídák’s post hoc analyses.

### Verteporfin induces dsDNA damage when combined with radiation and prevents repair of dsDNA breaks

We next investigated how verteporfin affects the DNA damage response to radiation in our chordoma models. Comet assays demonstrated that radiation induced dsDNA damage in chordoma cells in a dose-dependent manner, with 4 Gy of radiation required to significantly increase dsDNA damage (supplement). Neither 2 Gy radiation nor verteporfin alone at doses as high as 5μM significantly increased the magnitude of dsDNA damage. For reference, the IC50 of JHC7 was 2.8μM. The combination of verteporfin with 2 Gy radiation significantly increased dsDNA damage in JHC7 (3mM: **p<0.0001**, 5mM: **p<0.0001**), U-CH2 (3mM: **p=0.0042**, 5mM: **p<0.0001**), UM-Chor1 (5 mM: **p<0.0001**), and MUG-Chor1 (3mM: **p=0.0009**, 5mM: **p<0.0001**) (**Figure 2D**).

We then performed a series of experiments and found that while chordoma cells repair dsDNA damage over time, verteporfin inhibits the repair of dsDNA breaks. **Figure 2E** depicts the amount of dsDNA damage that develops over time after 2 Gy radiation and is graphed relative to a baseline value at 15 min after radiation. JHC7 and U-CH1 cells were able to repair DNA damage over time, but verteporfin prevented the repair of DNA damage with significantly increased dsDNA damage at 1 h (JHC7: **p<0.0001**, U-CH1: **p=0.0003**), 6 h (JHC7: **p<0.0001**, U-CH1: **p<0.0001**), 24 h (JHC7: **p=0.0059**, U-CH1: **p<0.0001**), 48 h (JHC7: **p<0.0001**, U-CH1: **p<0.0001**), and 96 h (JHC7: **p<0.0001**, U-CH1: **p<0.0001**).

### Verteporfin inhibits DNA damage repair proteins and genes

We next performed western blotting and qPCR assays to measure the ability of verteporfin to alter the expression of proteins and genes known to regulate the DNA damage repair cascade. Western blot analysis revealed that increasing amounts of verteporfin combined with radiation significantly decreased the expression of ATM, ATR, c-ABL, PARP, YAP, and pYAP (**Table 1**). qPCR experiments demonstrated that verteporfin significantly reduced the expression of ATM, ATR, PARP, RPA1, RAD51, cyclin D1, BRCA1, BRCA2, and PPP2R4 (**Table 2**). These results are shown in **Figure 3A-B**.

**Table 1.** Western Blot displaying expression of radiation related proteins after treatment with verteporfin and radiation.

| PROTEIN | 0GY 3UM | 0GY 5UM | 4GY 3UM | 4GY 5UM |
| --- | --- | --- | --- | --- |
| ATM | p=0.0447 | p=0.0195 | p=0.0280 | p=0.0012 |
| ATR | p=0.0052 | p=0.0044 | p=0.0021 | p=0.0007 |
| PARP | p=0.0014 | p=0.0005 | p=0.0004 | p=0.0001 |
| C-ABL | p<0.0001 | p<0.0001 | p<0.0001 | p<0.0001 |
| YAP | p<0.0001 | p<0.0001 | p<0.0001 | p<0.0001 |
| PYAP | p=0.0017 | p=0.0010 | p<0.0001 | p<0.0001 |
| YH2AX | p=0.1824 | p=0.1206 | p=0.0037 | p=0.0005 |

**Table 2.**
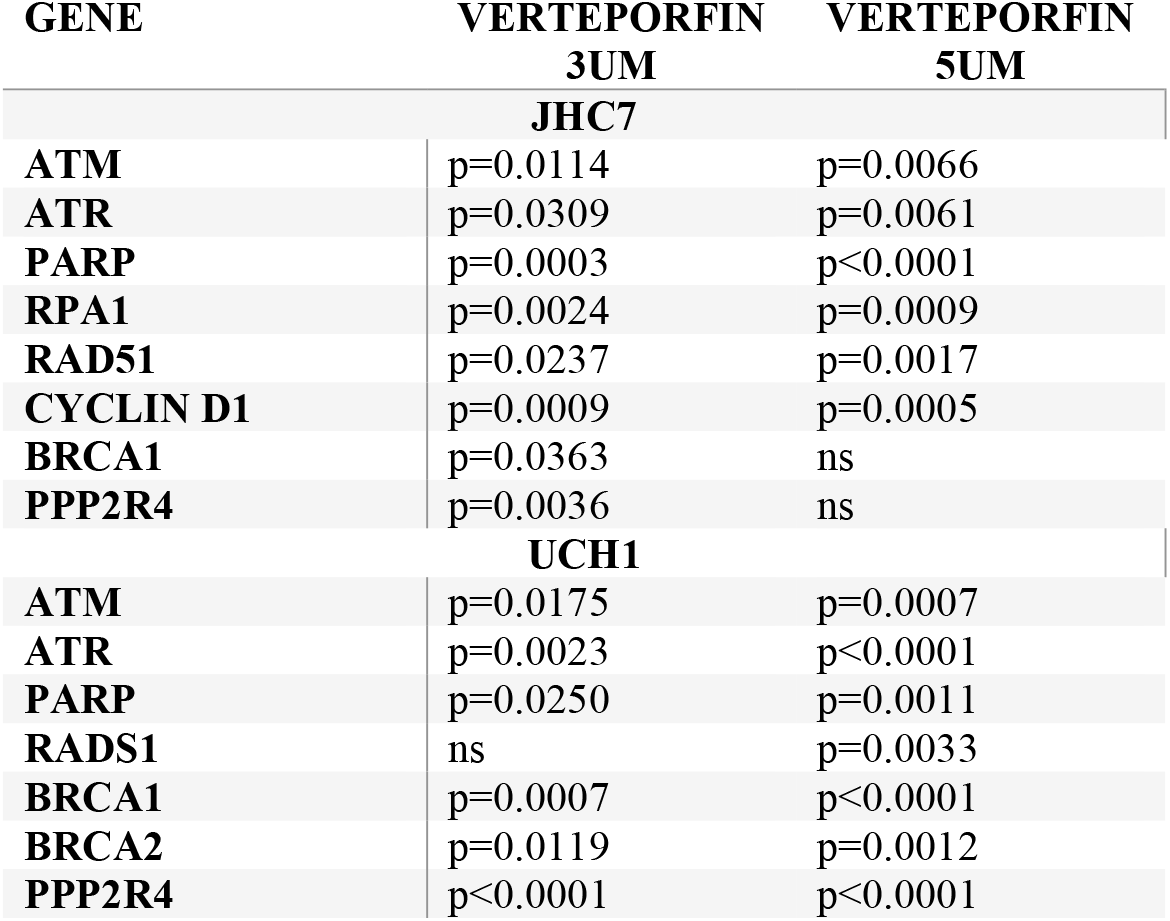
Quantitative PCR displaying expression of radiation related genes after treatment with verteporfin.

| GENE | VERTEPORFIN<br>3UM | VERTEPORFIN<br>5UM |
| --- | --- | --- |
| **JHC7** |  |  |
| ATM | p=0.0114 | p=0.0066 |
| ATR | p=0.0309 | p=0.0061 |
| PARP | p=0.0003 | p<0.0001 |
| RPA1 | p=0.0024 | p=0.0009 |
| RAD51 | p=0.0237 | p=0.0017 |
| CYCLIN D1 | p=0.0009 | p=0.0005 |
| BRCA1 | p=0.0363 | ns |
| PPP2R4 | p=0.0036 | ns |
| **UCH1** |  |  |
| ATM | p=0.0175 | p=0.0007 |
| ATR | p=0.0023 | p<0.0001 |
| PARP | p=0.0250 | p=0.0011 |
| RADS1 | ns | p=0.0033 |
| BRCA1 | p=0.0007 | p<0.0001 |
| BRCA2 | p=0.0119 | p=0.0012 |
| PPP2R4 | p<0.0001 | p<0.0001 |

**Figure 3.**
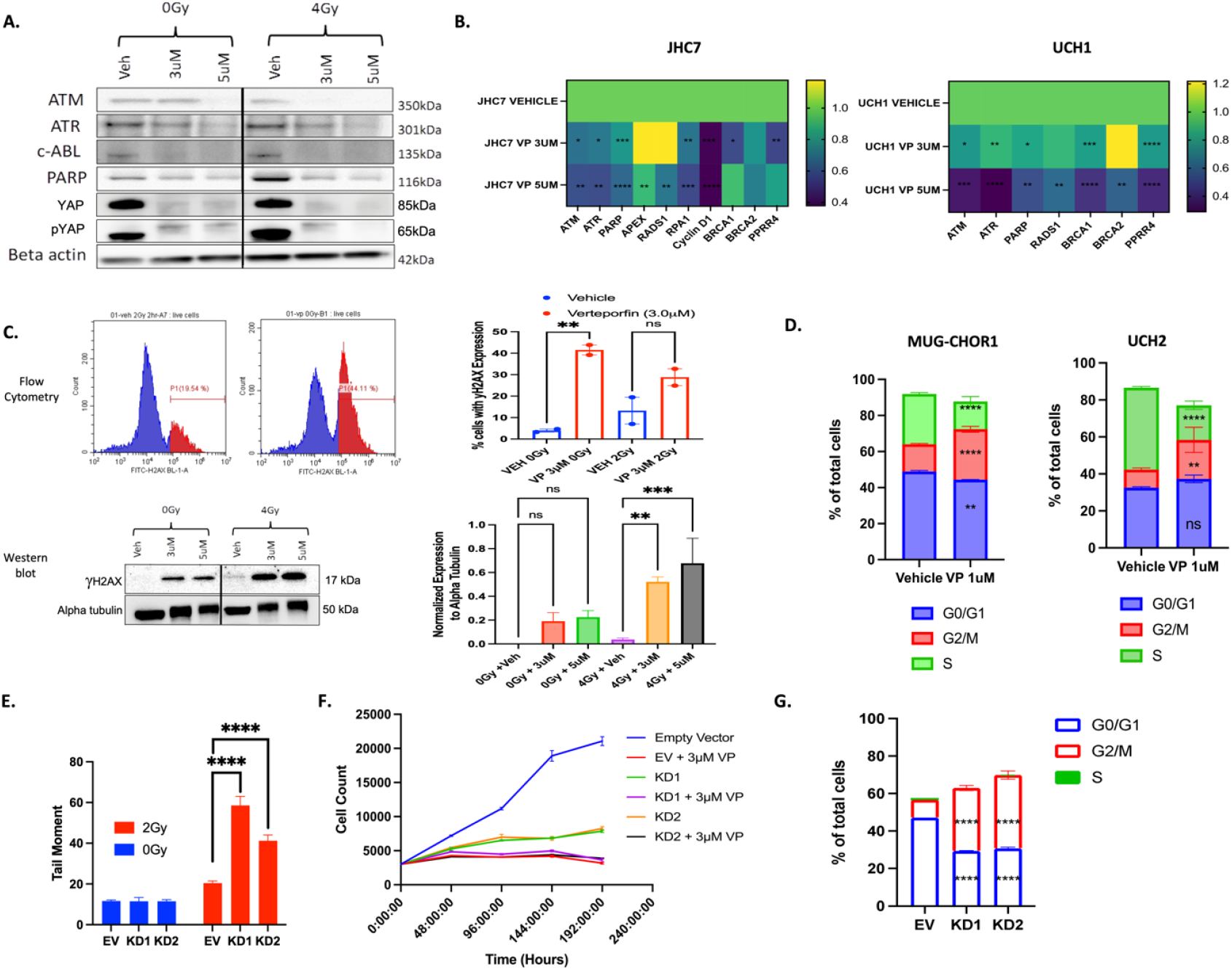
**A**. Verteporfin shuts down YAP and DNA damage repair proteins in JHC7 cells. Western blot revealing verteporfin significantly diminished protein expression of ATM (0Gy: 3μM **p=0.0447**, 5μM **p=0.0195**; 4Gy: 3μM **p=0.0280**, 5μM **p=0.0012**), ATR (0Gy: 3μM: **p=0.0052**, 5μM: **p=0.0044**; 4Gy: 3μM: **p=0.0021**, 5μM: **p=0.0007**), c-ABL (0Gy: 3μM **p<0.0001**, 5μM **p<0.0001**; 4Gy: 3μM **p<0.0001**, 5μM **p<0.0001**), PARP (0Gy: 3μM **p=0.0014**, 5μM **p=0.0005**; 4Gy: 3μM **p=0.0004**, 5μM **p=0.0001)**, YAP (0Gy: 3μM **p<0.0001**, 5μM **p<0.0001**; 4Gy: 3μM **p<0.0001**, 5μM **p<0.0001**). **B**. Verteporfin shuts down DNA damage repair genes. Heatmap signatures displaying significantly diminished gene expression in JHC7 and UCH1 cells following verteporfin treatment. There is significantly diminished gene expression of ATM (JHC7 (3μM: **p=0.0114**, 5μM: **p=0.0066**), UCH1 (3μM: **p=0.0175**, 5μM: **p=0.0007**)), ATR: JHC7 ((3μM: **p=0.0309**, 5μM: **p=0.0061**), UCH1 (3μM: **p=0.0023**, 5μM: **p<0.0001**), PARP: JHC7 (3μM: **p=0.0003**, 5μM: **p<0.0001**), UCH1 (3μM: **p=0.0250**, 5μM: **p=0.0011**), RADS1: JHC7 (5μM: **p=0.0017**), UCH1 (5μM: **p=0.0033**), RPA 1: JHC7 (3μM: **p=0.0024**, 5μM: **p=0.0009**), BRCA1: JHC7 (3μM: **p=0.0363**), UCH1 (3μM: **p=0.0007**, 5μM: **p<0.0001**), BRCA2: UCH1 (5μM: **p=0.0012**), PPP2R4: JHC7 (3μM: **p=0.0036**), UCH1 (3μM: **p<0.0001**, 5μM: **p<0.0001**). **C**. Verteporfin leads to increased γH2AX expression. Flow cytometry demonstrating increased γH2AX expression in JHC7 cells after verteporfin treatment (0GY: **p=0.0047**, 2Gy: p=0.0911). Western blot demonstrates significantly increased γH2AX expression (0Gy: 3μM p=0.1824, 5μM p=0.1206; 4Gy: 3μM **p=0.0037**, 5μM **p=0.0005**). **D**. Flow cytometry experiment demonstrating increased frequency of cells in G_2_M (MUG-CHOR1: **p<0.0001**, UCH2: **p=0021**) and decreased frequency of cells in S (MUG-CHOR1: **p<0.0001**, UCH2: **p<0.0001**) phases. **E**. YAP knockdown results in enhanced radiation induced DNA damage. Comet assays determined that YAP knockdown resulted in significantly enhanced double stranded DNA damage after exposure to 2Gy radiation (KD1: **p<0.0001**, KD2: **p<0.0001**). **F**. YAP Knockdown Decreases Cell Proliferation (Empty vector vs KD1: **p<0.0001**, empty vector vs KD2: **p<0.0001**). **G**. YAP knockdown results in increased frequency of cells in G2M phase (KD1 & KD2: **p<0.0001**). A-E, G: One-way and two-way ANOVA with Šídák’s post-hoc multiple comparisons test, F: Non-linear fit with extra sum-of-squares F test.

### Verteporfin increases gH2AX expression in chordoma cells

Next, we used flow cytometry and immunoblotting to measure gH2AX levels, an indirect measure of DNA damage (**Figure 3C**). We found a significant increase in gH2AX expression in JHC7 cells pretreated with verteporfin compared with vehicle treatment (0GY: **p=0.0047**, 2 Gy: p=0.0911). Western blotting also revealed a significant increase in gH2AX protein expression (4 Gy: 3μM **p=0.0037**, 5μM **p=0.0005**).

### Verteporfin increases the frequency of cells in G_*2*_M

The cell cycle changes that occurred secondary to verteporfin treatment of chordoma cell lines were determined by flow cytometry (**Figure 3D**). Verteporfin increased the relative number of cells in the G_2_M phase (UCH2: **p=0.0021**) and decreased the relative number of cells in the S (UM-CHOR1, **p<0.0001**; UCH2, **p<0.0001**) and G_0_/G_1_ (UM-CHOR1: **p=0.0042**) phases.

### YAP knockdown results in diminished cell growth, enhanced radiation sensitivity, and cell cycle arrest

To investigate whether our results were secondary to the YAP-mediated effects of verteporfin, we generated two YAP knockdown cell lines (KD1 and KD2) and performed a series of experiments with an empty vector (EV) control. Comet assays showed that YAP knockdown resulted in significantly increased dsDNA damage (KD1, **p<0.0001**; KD2, **p<0.0001**) after exposure to 2 Gy radiation (**Figure 3E**). YAP knockdown also led to significantly diminished cell proliferation (empty vector vs. KD1: **p<0.0001**, empty vector vs. KD2: **p<0.0001**), and the addition of 3mM verteporfin did not significantly diminish cell growth compared to YAP knockdown (**Figure 3F**). YAP knockdown also led to a significantly increased frequency of cells in G_2_/M (**p<0.0001**) and a significantly decreased frequency of cells in the G_0_/G_1_ (**p<0.0001**) cell cycle phase (**Figure 3G**).

### Verteporfin significantly decreases tumor growth in vivo

Given that verteporfin sensitizes chordoma cells to radiotherapy *in vitro*, we assessed its efficacy *in vivo* using a subcutaneous JHC7 chordoma xenograft model. Both systemically administered verteporfin (**p<0.0001**) and intratumoral administration of verteporfin-loaded microparticles (**p<0.0001**) significantly decreased tumor growth compared to that in the vehicle group (**Figure 4A**). We then evaluated the mice that received radiation and compared them with those that did not receive radiation. Our results demonstrate that the addition of radiation to verteporfin has an additional benefit over verteporfin alone. At 14 days post-treatment, there was significantly less tumor growth in the tumors that received radiation after treatment with verteporfin-loaded microparticles when compared to those that received verteporfin-loaded microparticles without radiation (**p=0.0098**) (**Figure 4B)**. The same was not true for the systemic verteporfin group and the control group, indicating that intratumoral verteporfin-loaded microparticles can sensitize tumors to radiation. Immunohistochemical analyses demonstrated that there was significantly diminished cytoplasmic (**p<0.05**) and nuclear (**p<0.0001**) YAP expression in tumors treated with verteporfin-loaded microparticles compared to the vehicle group (**Figure 4C**).

**Figure 4.**
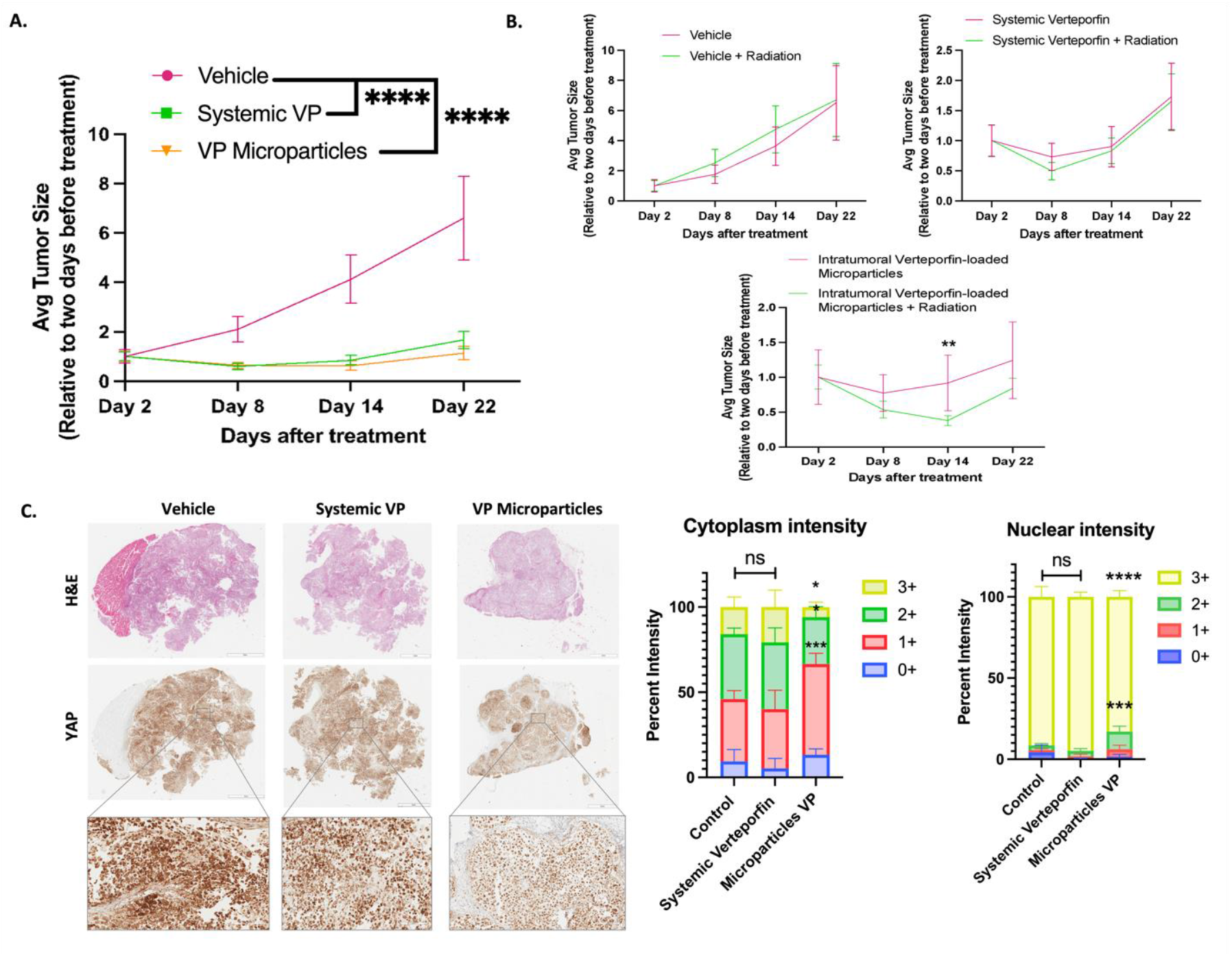
**A**. Verteporfin significantly slows the growth of tumors in vivo, both as verteporfin systemic drug (**p<0.0001**) and local intratumoral microparticle-loaded drug (**p<0.0001**). **B**. The addition of radiation significantly increased the tumor slowing effect of intratumoral verteporfin-loaded microparticles at 14 days post-treatment (**p=0.0098**). **C**. Immunohistochemistry percentage of cytoplasmic and nuclear YAP intensity was measured using the Aperio ImageScope x64 Software (Aperio Systems). Signal intensity ranged from 0+ (being the lowest detected signal) to 3+ (determined as the strongest detected signal). There was significantly less 1+ (**p<0.002**) 2+ (**p<0.05**) and 3+ (**p<0.05**) percentage intensity between the control and the verteporfin microparticles group. The same was true for nuclear YAP intensity, in which there was significantly less 2+ (**p<0.005**) and 3+ (**p<0.0001**) percentage intensity. Two-way ANOVA followed by a Tukey’s multiple comparisons test.

## DISCUSSION

Our results demonstrate that inhibiting YAP sensitizes chordoma to radiation, enhancing tumor control at lower and less toxic doses of radiation. We discovered two mechanisms that drive this effect: 1) inhibition of YAP inhibits the DNA damage repair cascade and 2) inhibition of YAP increases the frequency of cells in G_2_M, where they are more sensitive to radiation. Verteporfin is an FDA-approved drug that is a potent YAP inhibitor. We found that verteporfin inhibits the growth of chordoma cells and sensitizes these tumors to radiation. YAP knockdown demonstrated similar results, indicating that the effect of verteporfin was targeted. Our verteporfin-loaded microparticles were effective *in vivo*, inhibiting the growth of chordoma tumors and making them more sensitive to radiation. Verteporfin has a half-life of 6 h; therefore, the encapsulated formulation of this drug with slow release would be expected to significantly prolong therapeutic drug concentrations. This is particularly important for a slowly growing tumor, such as chordoma, because there is a very low probability that any tumor cell will be in G_2_M at any given time, which increases radioresistance. Prolonging the therapeutic concentrations of verteporfin is expected to increase the fraction of tumor cells in G_2_M, thereby enhancing radiosensitivity.

Phosphorylation of H2AX to generate gH2AX is the first step in the DNA damage cascade and is the earliest cellular response to radiation, upregulated just seconds after radiation exposure. ^23^ It has been used as a measure of the DNA damage response. ^24,25^ Hao *et al* used changes in gH2AX as a surrogate marker of DNA damage to demonstrate that protein phosphatase 2A (PP2A) enhanced DNA damage in chordoma. ^26^ We also found that gH2AX significantly increased after radiation in chordoma cells pretreated with verteporfin. In addition to indirectly measuring DNA damage using gH2AX, we directly measured DNA damage using comet assays and found that both verteporfin and YAP knockdown caused a significant increase in dsDNA breakage after ionizing radiation. To our knowledge, this is the first time that this has been observed in chordomas.

Our study also provides insight into the mechanisms by which radiosensitization occurs. ATM and ATR are two of the main effectors of the DNA damage cascade. Verteporfin inhibits the expression of ATM/ATR and has either a direct or indirect effect on many downstream effectors in the DNA damage cascade, including RAD51, PARP, APEX, RPA1, and BRCA. Cells are most sensitive to radiation in the G_2_M phase of the cell cycle, and we found that verteporfin and YAP knockdown causes G_2_M arrest, leading to an accumulation of cells that are more sensitive to damage from ionizing radiation.

Recent data indicate that YAP is involved in radiation resistance in various tumors. Li et al. found that YAP facilitates radiation resistance in nasopharyngeal carcinoma secondary to the nuclear translocation of YAP. ^27^ Similar data have been published for breast, brain, colorectal, and endometrial cancers ^28,29-32^ showing that YAP translocation into the nucleus is the first step in radiation resistance in these tumors, and our study provides insight into what happens after YAP enters the nucleus.

This study has significant clinical implications that require further exploration. It is important to note that neither 2 Gy radiation nor 5 mM verteporfin alone significantly increased dsDNA damage in chordoma cells, but the combination of the two doses. This highlights a potential therapeutic strategy for patients with chordoma, especially for those with recurrent disease. Recurrent tumors are known to be more resistant to therapy, especially those with prior radiation. Verteporfin-loaded microparticles can be injected directly into the tumor via a percutaneous method, followed by radiation treatment, providing a nonsurgical alternative treatment for these patients. Our data show that YAP inhibition also inhibits the growth of these tumors; therefore, there would be a two-fold benefit for these patients. This aspect needs to be explored in future clinical trials.

## CONCLUSIONS

In this study, we determined that inhibition of the YAP regulatory axis decreased growth and enhanced the radiosensitivity of chordoma tumors. This can be achieved with verteporfin using novel verteporfin-loaded microparticles. Future orthotopic animal models are needed to further translate this into human clinical trials so that one day, we can possibly have more therapies in our armamentarium for this devastating cancer.

## Funding

O.O.A. was funded by a Neurosurgery Research and Education Foundation (NREF) grant.

## Conflicts of Interest

Authors declare no conflicts of interest.

## Data Availability

The data will be made available upon reasonable request.

## Declaration of Author Contributions

OOA: performed experiments, analyzed data, wrote the manuscript, created figures/tables, confirmed the final version of the manuscript; PSM: performed experiments, analyzed data, manuscript writing, critically revised manuscript, created figures; PS: edited figures, analyzed data, performed experiments, critically revised manuscript; RW: analyzed data, performed experiments; SYT: performed experiments, critically revised the manuscript; SSR: critically revised manuscript; SK: critically revised manuscript; CP: critically revised manuscript; ZLG: critically revised manuscript; JJG: critically revised manuscript; AQH: study supervision, critically revised manuscript. All authors have reviewed the final version of the manuscript.

## Notes

### Competing Interest Statement

The authors have declared no competing interest.

