## Supplementary material for "Inhibition of Yes-Associated Protein (YAP) with Verteporfin Enhances Radiosensitivity in Chordoma by Inducing G_2_M Arrest and Inhibiting the DNA Damage Response": N/A

| Supplementary Table 1. Primer Sequences | | |
|---|---|---|
| ATM | ATM F | TTGATCTTGTGCCTTGGCTAC |
| | ATM R | TATGGTGTACGTTCCCCATGT |
| ATR | ATR F | TCCCTTGAATACAGTGGCCTA |
| | ATR R | TCCTTGAAAGTACGGCAGTTC |
| PARP | PARP-1 F | CGGAGTCTTCGGATAAGCTCT |
| | PARP-1 R | TTTCCATCAAACATGGGCGAC |
| APEX | APEX F | CAATACTGGTCAGCTCCTTCG |
| | APEX R | TGCCGTAAGAAACTTTGAGTGG |
| RADS1 | RAD51 F | CAACCCATTTCACGGTTAGAGC |
| | RAD51 R | TTCTTTGGCGCATAGGCAACA |
| RPA1 | RPA1 F | GTGGACCATTTGTGCTCGTG |
| | RPA1 R | TTCGTCAACCAGTTCTAGGGA |
| CyclinD1 | Cyclin D1 F | GCTGCGAAGTGGAAACCATC |
| | cyclin D1 R | CCTCCTTCTGCACACATTTGAA |
| BRCA1 | BRCA1 F | GCTCGTGGAAGATTTCGGTGT |
| | BRCA1 R | TCATCAATCACGGACGTATCATC |
| BRCA2 | BRCA2 F | AGGATCTCTCACCGCAGCTAA |
| | BRCA2 R | TCAGGCCCAACATCAAGTGTG |
| PPRR4 | PPP2R4 F | TCTCAGGCATACGCTGACTAC |
| | PPP2R4 R | GGAGACTCTGTACTCGAAGGT |

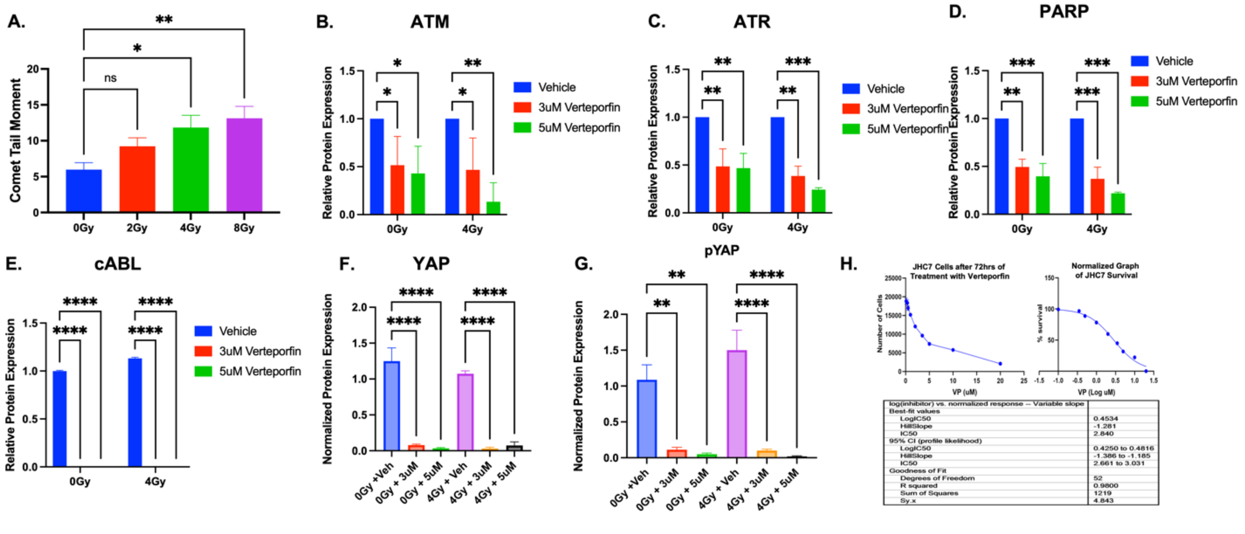


**Supplementary Figure 1**. A. Radiation induces double stranded breaks in JHC7 chordoma cells in a dose dependent manner (4Gy: **p=0.0127**, 8Gy: **p=0.0011**). B. Western blot relative protein expression for ATM shows diminished protein expression after verteporfin treatment: (0Gy: 3μM **p=0.0447**, 5μM **p=0.0195**; 4Gy: 3μM **p=0.0280**, 5μM **p=0.0012**). C. Western blot relative protein expression for ATR shows diminished protein expression after verteporfin treatment: (0Gy: 3μM: **p=0.0052**, 5μM: **p=0.0044**; 4Gy: 3μM: **p=0.0021**, 5μM: **p=0.0007**). D. Western blot relative protein expression for PARP shows diminished protein expression after verteporfin treatment: PARP (0Gy: 3μM **p=0.0014**, 5μM **p=0.0005**; 4Gy: 3μM **p=0.0004**, 5μM **p=0.0001).** E. Western blot relative protein expression for c-ABL shows diminished protein expression after verteporfin treatment: c-ABL (0Gy: 3μM **p<0.0001**, 5μM **p<0.0001**; 4Gy: 3μM **p<0.0001**, 5μM **p<0.0001).** F. Western blot relative protein expression for YAP shows diminished protein expression after verteporfin treatment: γH2AX (0Gy: 3μM **p<0.0001**, 5μM **p<0.0001**; 4Gy: 3μM **p<0.0001**, 5μM **p<0.0001)**. G. Western blot relative protein expression for YAP shows diminished protein expression after verteporfin treatment: pYAP (0Gy: 3μM **p=**0.0017, 5μM **p=**0.0010; 4Gy: 3μM **p<0.0001**, 5μM **p<0.0001)**. H. Verteporfin IC50 of JHC7 chordoma cells is 2.8uM.
